# Aborting meiosis overcomes hybrid sterility

**DOI:** 10.1101/2020.12.04.411579

**Authors:** Simone Mozzachiodi, Lorenzo Tattini, Agnes Llored, Agurtzane Irizar, Neža Škofljanc, Melania D’Angiolo, Matteo De Chiara, Benjamin P. Barré, Jia-Xing Yue, Angela Lutazi, Sophie Loeillet, Raphaelle Laureau, Souhir Marsit, Simon Stenberg, Benoit Albaud, Karl Persson, Jean-Luc Legras, Sylvie Dequin, Jonas Warringer, Alain Nicolas, Gianni Liti

## Abstract

Hybrids between species or diverged lineages contain fundamentally novel genetic combinations but an impaired meiosis often makes them evolutionary dead ends. Here, we explored to what extent and how an aborted meiosis followed by a return-to-growth (RTG) promotes recombination across a panel of 20 yeast diploid backgrounds with different genomic structures and levels of sterility. Genome analyses of 284 clones revealed that RTG promoted recombination and generated extensive regions of loss-of-heterozygosity in sterile hybrids with either a defective meiosis or a heavily rearranged karyotype, whereas RTG recombination was reduced by high sequence divergence between parental subgenomes. The RTG recombination preferentially occurred in regions with local sequence homology and in meiotic recombination hotspots. The loss-of-heterozygosity had a profound impact on sexual and asexual fitness, and enabled genetic mapping of phenotypic differences in sterile lineages where linkage or association analyses failed. We propose that RTG gives sterile hybrids access to a natural route for genome recombination and adaptation.

**One sentence summary:** Aborting meiosis followed by a return to mitotic growth promotes evolution by genome wide-recombination in sterile yeast hybrids.

## Main text

Meiotic recombination is a primary source of genetic diversity in species undergoing sexual reproduction. During meiosis, crossovers ensure that haploid gametes receive one copy of each chromosome and prevent a genetic imbalance in the offspring that often decreases fitness (*1*). However, reproductive barriers acting before (pre-zygotic) or after (post-zygotic) zygote formation (*2*) can arise during species evolution, with post-zygotic barriers leading to hybrid sterility (*3*) across the tree of life (*4*).

*Saccharomyces* hybrids have served as models to elucidate the mechanisms contributing to post-zygotic reproductive isolation (*5*). Sterile *Saccharomyces* intraspecies and interspecies hybrids have been repeatedly isolated in both wild and domesticated environments (*6*). Relaxed selection on sexual reproduction in domesticated *S. cerevisiae* populations has led to the accumulation of loss-of-function mutations in genes involved in gametogenesis, i.e. “sporulation” in yeast biology, and to severe sterility (*7*). Chromosomal rearrangements between the subgenomes of a hybrid can also lead to sterility due to aberrant chromosome pairing and segregation (*8, 9*). The Malaysian *S. cerevisiae* lineage represents the most dramatic chromosomal speciation example as it contains 5 rearranged chromosomes that isolate it reproductively from other *S. cerevisiae* lineages, despite retaining high levels of sequence similarity (*10, 11*). In contrast, *Saccharomyces* interspecies hybrids produce inviable gametes because of the extreme DNA sequence divergence between parental subgenomes, namely heterozygosity, which suppresses recombination and leads to chromosome mis-segregation (*12*). Therefore, many *Saccharomyces* intraspecies and interspecies hybrids have only very limited possibilities to evolve through meiosis and some are completely sterile. Moreover, wild yeasts reproduce sexually approximately only once every 1000 asexual generations, thus further limiting the role of meiosis in both intraspecies and interspecies evolution (*13*).

We showed that aborting the meiosis in a fertile *S. cerevisiae* lab hybrid and returning the diploid cells to mitotic growth, a process known as “return-to-growth” (RTG) (*14*), reshuffles the diploid genome and produces extensive regions of loss-of-heterozygosity (LOH) (*15*). LOHs resulting from single crossing-over events generate terminal events that extend to the chromosome ends, whereas interstitial LOHs in chromosome cores can be ascribed to gene-conversions or double crossovers. Meiosis in *S. cerevisiae* is initiated by starvation and chromosomes are replicated during the meiotic S-phase (**Fig. 1A**). Next, as in most eukaryotes (*16*), a DNA topoisomerase, Spo11p in yeast, generates genome-wide double-strand breaks (DSBs) (*17*). The repair of these DSBs produces joint DNA molecules and ultimately leads to recombination through chromosomal crossover (CO) and non-crossover (NCO) molecules. Yeast cells brought back to a nutrient-rich environment before the commitment to complete meiosis, abort the meiotic program and instead express genes promoting mitotic division, thus entering the RTG process (*18, 19*). RTG cells repair the DSBs, bud likewise mitotically dividing cells, and segregate their recombined chromatids with no further DNA replication (*20*). This process generates a mother and a daughter cell that maintain the parental diploid state, as they do not complete the two chromosomal segregation events that occur in a normal meiosis, but contain different rearranged genotypes. Here, we show that RTG allows sterile *Saccharomyces* hybrids to overcome common reproductive barriers, and we propose RTG as a powerful alternative recombination mechanism that may play an unanticipated role in the evolution of hybrid genomes in nature.

**Figure 1.**
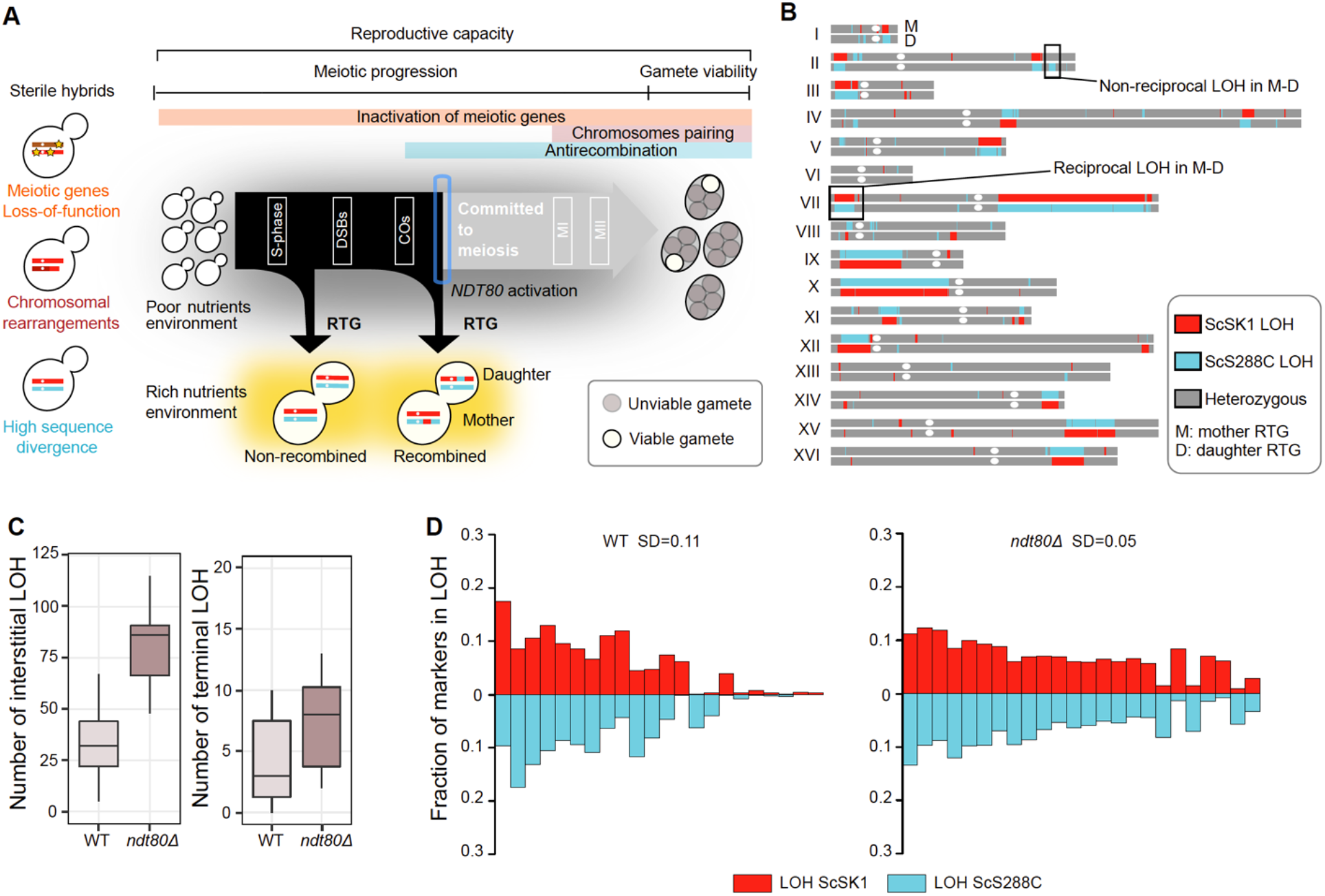
RTG paradigm and genomic landscape of *ndt80Δ* evolved RTG clones. (**A**) Inactivation of essential meiotic genes (yellow stars), genomic rearrangements and high levels of heterozygosity in hybrid genomes can impair meiotic progression or reduce gamete viability. RTG represents an alternative route to hybrid evolution that creates LOH blocks. (**B**) Example of LOH map in a mother (top)-daughter (bottom) RTG pair derived from the *ndt80Δ* hybrid. LOH blocks can be tracked by genotyping single-nucleotide marker in the two parental subgenomes, which give a homozygote readout if the position is in a LOH block. (**C**) Boxplots of the number of interstitial (left) and terminal LOH events (right) in the WT (*n*=22) and *ndt80Δ* (*n*=24) RTG clones derived from ScS288C/ScSK1. *ndt80Δ* RTGs have more interstitial (*p*-value=3.9×10^−8^) and terminal (*p*-value=0.001) LOHs. Both tests are one-tailed Wilcoxon rank-sum test, with continuity correction. We performed the same statistical test by taking into account only the mother cell in both datasets (*n=*12 *ndt80Δ, n=*11 WT) and confirmed that *ndt80Δ* mothers accumulated more interstitial (*p-*value=5.033×10^−8^) and terminal (*p-*value=0.0053) LOHs than the WT mothers. No LOH parental bias was detected by comparing the average number of events towards one of the two subgenomes in WT (Welch’s t-Test, *p*-value = 0.67) and *ndt80Δ* datasets (Welch’s t-Test, *p*-value = 0.25). (**D**) Fraction of homozygous markers for both parental alleles for the WT and *ndt80Δ* datasets. Each bar represents a sequenced clone. The *ndt80Δ* RTGs show a lower LOH variation compared to the wild-type, both if considered as a pooled dataset (*n=*24) (Fligner-Killeen test, *p*-value = 0.0008) and if considering only the mothers from each datasets (Fligner-Killeen test, *p-*value=0.02).

### An incomplete meiosis supports recombination in a sterile hybrid

Deleterious variants in meiotic genes can lead to various meiotic defects and sterility. However, mutations that impair mid or late meiotic progression should not prevent RTG, if cells can resume mitotic-like chromosome segregation and budding (*20, 21*) (**Fig. 1A**). Therefore, we probed whether RTG allows *S. cerevisiae* (Sc) to overcome mutational barriers in meiosis by deleting *NDT80*, the master regulator of middle and late meiotic genes, in an otherwise fertile ScS288C/ScSK1 intraspecies lab-hybrid. First, we confirmed by DAPI staining that *ndt80Δ* cells were arrested before the first meiotic division (MI) after 8 hours of sporulation induction, whereas ≥50% of the wild-type ScS288C/ScSK1 cells had already completed MI (≥ 2 nuclei). Then, we evolved the hybrid through the RTG protocol and sequenced the genome of 12 mother-daughter *ndt80Δ* RTG pairs (*n*=24) isolated 8 hours after sporulation induction. As a comparison, we re-analysed the mother-daughter RTG genomes (*n=*22) of the wild-type ScS288C/ScSK1 derived from 4-5 hours of sporulation induction (*15*). Short-read sequencing data were analysed by an integrated framework that enables a high-resolution view also of LOHs supported by only a single marker (*22*). The *ndt80Δ* clones had highly recombined diploid genomes (**Fig. 1B**) with an average of 88 LOHs per clone, when combining both terminal (*n*=181) and interstitial (*n=*1933) LOHs. The number of interstitial LOHs was higher in the *ndt80Δ* clones (**Fig. 1C, S1A**), which also had more markers lying in non-reciprocal recombined regions (*p*-value = 3.663^-06^, Welch’s t-test) (**Table S13**). Consistently, the *ndt80Δ* background accumulated more NCOs (*23*). In contrast, fewer large LOHs (> 10 kbp) were detected compared to the wild-type (**Fig. S1A, S2A**). Despite the LOH size variability, the two genetic backgrounds showed a similar median fraction of markers in LOH (**Table S6**), but the distribution was more homogeneous in the *ndt80Δ* population compared to the wild-type population (**Fig. 1D**), in line with cells accumulating at the same stage of meiosis by arresting at prophase I. The frequency of LOH events increased from the centromere towards chromosome ends in both datasets, in accordance with centromeres being cold meiotic recombination regions (**Fig. S1C**). Finally, we found no bias in LOH formation towards either of the two parental subgenomes in the datasets, consistent with RTG generating complementary recombined genomes with few non-reciprocal events (*15*) (**Fig. 1D**). Overall, we demonstrated that RTG allows a hybrid that is unable to complete meiosis due to the lack of a functional key meiotic gene to generate highly recombined diploid genomes.

### Profiling RTG efficiency across hybrid diversity

Next, we asked to what extent RTG promotes recombination in hybrids with variable degrees of sterility due to heterozygosity and structural differences between the subgenomes. We exploited natural variation to generate a panel of 19 diploid genetic backgrounds, comprised by 4 fully homozygous *S. cerevisiae* (Sc), 7 *S. cerevisiae* intraspecies hybrids (Sc/Sc) and 8 interspecies hybrids between *S. cerevisiae* and its sister species *S. paradoxus* (Sp/Sc) (**Fig. 2A-B, Table S1 and S9**). Then, we engineered a simple genetic system to measure RTG-induced LOH rates at the *LYS2* locus on chromosome II (**Fig. 2C**). We replaced one of the *LYS2* alleles with a *URA3* gene and compared how often this *URA3* marker was lost due to LOH, in cells returned to growth after 6 hours of meiosis induction (T6) but before commitment to meiosis of the fastest sporulating strain (**Fig. S3A**), to control mitotic cells (T0). We derived the efficiency of RTG-induced LOH (T6/T0) (**Fig. 2D and Tables S4**) and observed pronounced genetic background effects with fast sporulating strains promoting efficient recombination upon RTG (**Fig. 2E, Tables S2**). Among the homozygous diploid, the ScNA/ScNA showed the largest LOH rate increase (21-fold) (**Fig. 2D)** consistent with its faster meiotic progression, while the 3 other homozygous diploids had a poor sporulation efficiency and synchrony (**Fig. S3B**), preventing the production of a significant number of recombinant RTGs. All the intraspecies hybrids with a ScNA subgenome also showed a significant increase of recombination upon RTG, ranging from 3 to 42-fold, whereas other intraspecies hybrids showed no significant increase (**Fig. 2D**). In particular, we detected a 23-fold increase in LOH upon RTG in the ScMA/ScNA hybrid, showing that the non-collinearity between several homologous chromosomes other than chromosome II, did not reduce RTG-induced LOH at the *LYS2* locus.

**Figure 2.**
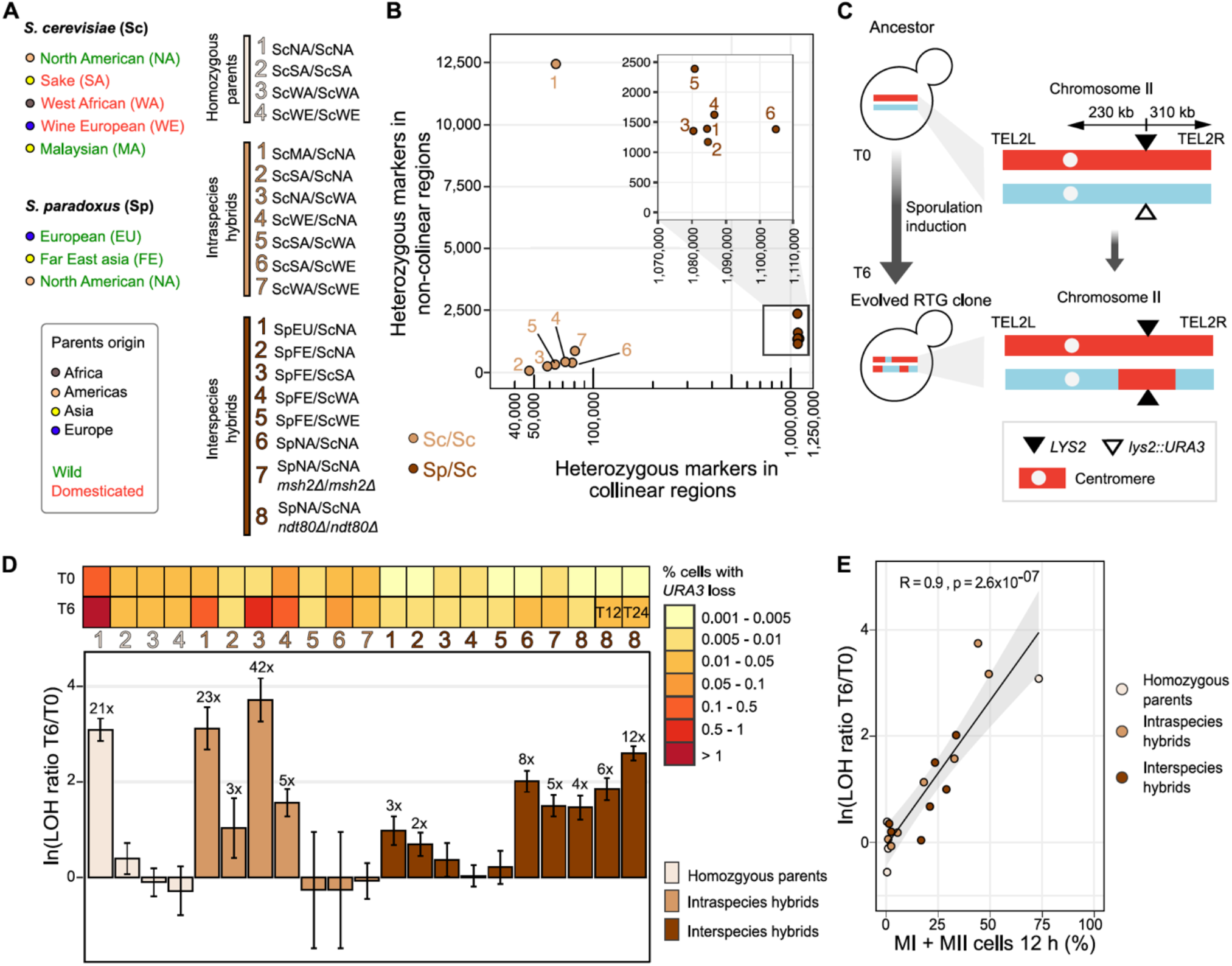
Quantifying RTG-induced recombination across hybrid diversity. **(A)** Geographical (coloured circle) and ecological (coloured name) origins of the parental strains (left) used for generating the homozygous diploids, intraspecies hybrids and interspecies hybrids (right). Diploids are grouped according to their level of heterozygosity as: homozygous parents (light brown), intraspecies hybrids (orange) and interspecies hybrids (dark brown) and the same colour code apply to panels B, E and E. **(B)** Level of heterozygosity across the hybrid panel with number of heterozygous markers detected in non-collinear and collinear regions. Each data point is labelled with a number/colour encoded according to the strains list on panel A. The four homozygous parents are not reported. **(C)** *URA3*-loss assay used for measuring RTG-induced recombination rates. **(D)** Top plot: percentage of cells growing on 5-FOA measured with the *URA3*-loss assay at two time points (or more, if specified): T0 = no meiosis induction, T6, T12 and T24 indicate 6, 12, and 24 hours of meiosis induction respectively. Each square is coloured accordingly to the average of the replicates as indicated in the right scale. The percentage of cells with *URA3*-loss was calculated as the ratio of colony forming units (CFU) CFU/mL in 5-FOA and CFU/mL in YPD. Bottom: bar plot showing the natural logarithm ratio of number of cells growing on 5-FOA at the RTG induction time (T6, or more) and control (T0). Error bars represent the standard deviation for each ratio. The number on the top of each bar plot indicates the linear fold increase (e.g. 24x) of the LOH rate, comparing T6 to T0 as specified above. **(E)** RTG efficiency as a function of meiotic progression after 12 hours measured as cells that passed first (MI) and second (MII) meiotic divisions. R is the Pearson’s correlation coefficient.

All the interspecies *S. cerevisiae/S. paradoxus* hybrids had a lower basal LOH rate during mitosis (10-fold average) and RTG enhanced the LOH rate less than in intraspecies hybrids (**Fig. 2D, Table S4**). Nevertheless, all three interspecies hybrids with a ScNA subgenome showed significant LOH rate increases (**Fig. 2D**). The lower LOH rates at both T0 and T6 suggest that very high heterozygosity strongly inhibits recombination in interspecies hybrids through an antirecombination mechanism, as it does during meiosis (*24*). To test this conjecture, we deleted the gene *MSH2*, which encodes a key protein involved in the mismatch machinery, in both subgenomes of the SpNA/ScNA hybrid. Indeed, we observed increased LOH rates in both T0 (3.6-fold) and T6 (2-fold) samples compared to the wild-type SpNA/ScNA hybrid (**Fig. 2D, Table S4**), despite meiotic progression being slower in the *msh2Δ* mutant (**Fig. 2E and S3B**). This is consistent with the mismatch repair machinery acting to prevent RTG recombination in hybrids with highly diverged subgenomes.

Finally, we generated a SpNA/ScNA *ndt80Δ* with both high heterozygosity and meiotic progression defects, and we measured LOH rates at 6 (T6), 12 (T12) and 24 (T24) hours after sporulation induction. We observed that RTG recombination increased with the time spent in gametogenesis, consistently with the fraction of cells engaged in RTG increasing with time (**Fig. 2D**). Hence, genetically very similar cells in clonal populations are nevertheless quite heterogeneous in their meiotic progression, with only a minor fraction of cells having committed to recombination at 6 hours. Thus, the absence of RTG induced recombination in a hybrid may reflect that cells had not progressed sufficiently in their meiosis to engage in recombination and produce recombinant RTGs. We therefore focused our follow-up whole-genome-sequencing analyses on hybrids for which the RTG did increase recombination. Overall, our experimental system showed that RTG efficiency varies broadly across genetic backgrounds and is strongly influenced by their meiotic progression.

### RTG recombination in hybrids with extensive chromosomal rearrangements

We performed whole-genome sequencing of RTG-evolved clones derived from the fertile ScWE/ScNA hybrid (*n=*24, T6) and the sterile ScMA/ScNA hybrid (*n=*123 T6, *n=*2 T4, *n=*3 T0), which have similar levels of heterozygosity but different genome structures (**Fig. 2B and Table S9**). The ScMA/ScNA hybrid represents sterility driven by chromosomal rearrangements, with 5 homologous chromosomes for which the parental subgenomes are not collinear over long stretches. Our analyses revealed that the RTG genomes of ScWE/ScNA and ScMA/ScNA clones recombined equally often (average number of LOH per clone=65 and 69, respectively; **Fig. 3A-B and Table S6**). This underscores that genome-wide RTG recombination is not hampered by extensive non-collinearity between subgenomes, in line with that the sterility caused missegregation of the non-collinear chromosomes rather than a lack of recombination (*25*). Moreover, neither parental subgenome was favoured over the other in terms of the created homozygosity (**Fig. S2C**) consistent with no parental bias in the accumulation of Spo11p-induced DSBs, which occur at highly conserved sites across the *S. cerevisiae* and *S. paradoxus* lineages used here (*26*).

**Figure 3.**
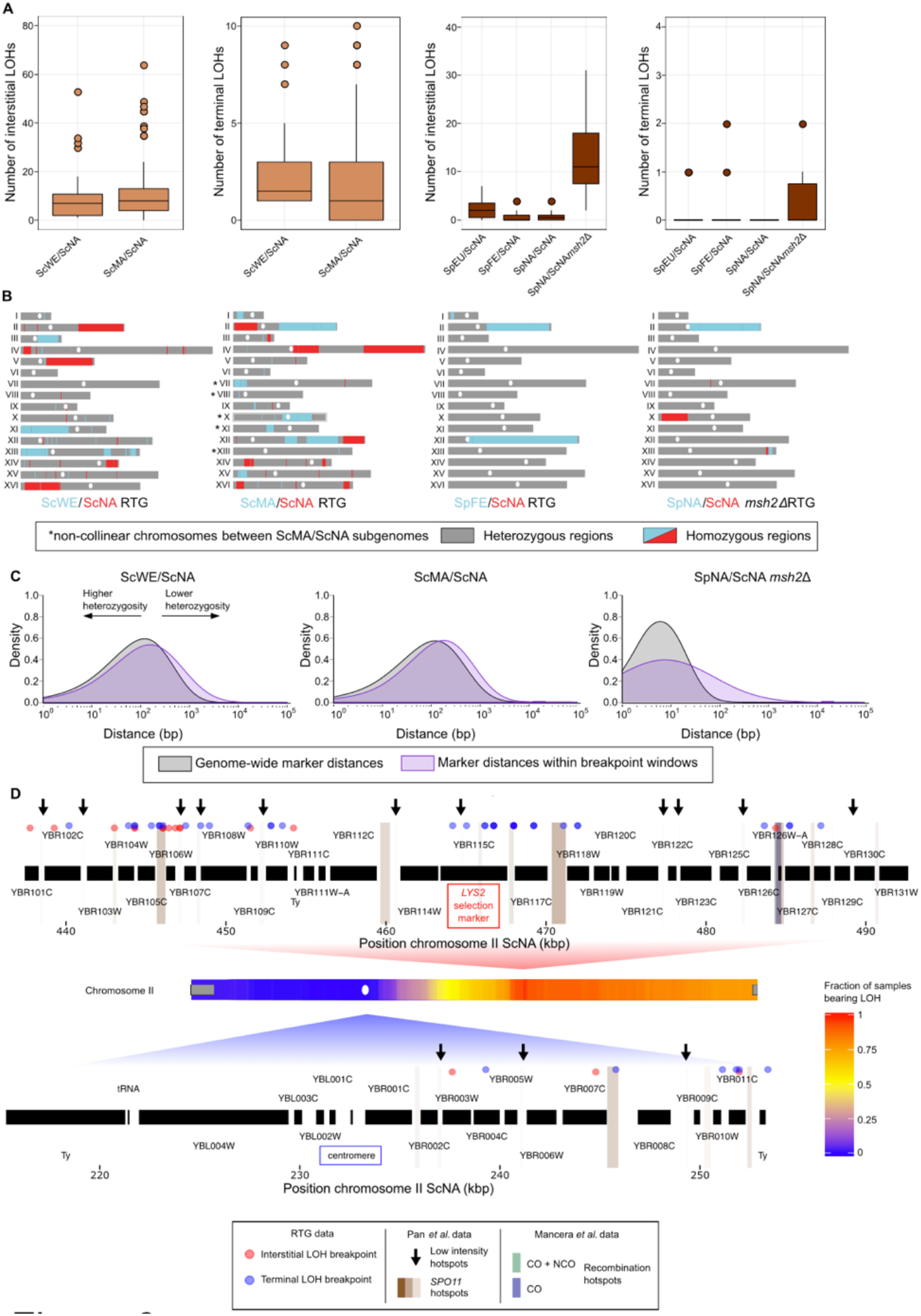
LOH landscape of hybrids evolved through RTG. (**A**) Left panels: boxplots of the number of interstitial and terminal LOHs in intraspecies hybrids. We detected no significant difference in the number of interstitial or terminal events comparing the ScWE/ScNA and the ScMA/ScNA datasets (two-sided Wilcoxon rank-sum test, *p*-value = 0.5 and *p*-value = 0.1, respectively). Right panels: boxplots of the number of interstitial and terminal LOHs in interspecies hybrids. Colour code as in figure 2A. (**B**) Genome-wide view of the LOH landscape for two RTG-evolved intraspecies hybrids (left) and two evolved interspecies hybrids (right). All plots are based on the ScNA reference genome. (**C**) Distribution of marker distances genome-wide (grey) and in LOH breakpoint windows (purple) across three different hybrids. LOH breakpoint windows comprise the 5 heterozygous markers and the 5 homozygous markers closer to the breakpoint. (**D**) Association between meiotic recombination and LOH breakpoints within two regions of chromosome II. Low-intensity Spo11p hotspots are highlighted with black arrows. The heatmap of the LOHs (i.e. two copies of ScNA alleles) resulting from the breakpoints is also reported. Grey boxes represent subtelomeric regions.

Both hybrids showed that the genomic intervals encompassing LOH breakpoints on the right arm of chromosome II were characterised by lower heterozygosity (**Fig. 3C, Fig. S4A-B)**. Moreover, LOH breakpoints on the right arm of chromosome II were associated both with known meiotic hotspots (*27*) and sites where Spo11p induces DSBs during meiosis (*28*) (**Fig. 3D, Fig. S4C, Table S10)**. Since this implies that RTG recombination relies on DSBs created during the aborted meiosis, we probed whether LOH breakpoints induced by RTG recombination were associated with meiotic recombination hotspots genome-wide and found this to be the case (**Fig. S4C, Table S11)**. We found that centromeres were always maintained in a heterozygous state, consistent with mitotic like segregation in RTG, and constituted RTG-induced recombination coldspots **(Fig. S1C)**. We found no increase in single nucleotide variants (SNVs), aneuploidies or copy number variations (CNVs) and conclude that RTG-induced recombination does not cause global genome instability (**Fig. S5A**). This is consistent with the major role of the error-free homologous recombination pathway in repairing meiotic DSBs.

Four ScWE/ScNA and nine ScMA/ScNA RTG clones carried a much larger fraction of markers in LOH (15-34% and 14-35% respectively) than the median per RTG clone (1% and 1.7%, respectively) (**Fig. S2C**). We hypothesized that extensive LOH could homogenise subgenomes sufficiently to alleviate the sterility of the ScMA/ScNA hybrid. Thus, we probed the capacity of three ScMA/ScNA RTG clones with extensive LOH to produce viable gametes. Indeed, these clones had up to 3-fold increased gamete viability compared to the ancestral ScMA/ScNA hybrid (**Fig. S6A and Table S3**), confirming that RTG can help restore meiotic fertility by homogenising subgenomes and making them more collinear.

We also detected LOHs encompassing the *MAT* locus on chromosome III in 2 of the ScWE/ScNA and in 1 of the ScMA/ScNA extensively recombined clones (**Fig. S6B**). We found that although this *MAT* homozygosity led to complete sterility and prevented further RTG cycles, it enabled these hybrids to pass from a diploid mating deficient behaviour to a haploid-like mating proficient, thereby providing a direct route to polyploidization (**Fig. S6 C-D**). Overall, these results showed that sterile intraspecies hybrids can evolve through RTG and overcome reproductive barriers caused by a divergent chromosomal structure, with even a single RTG cycle having a dramatic impact on the genome evolution.

### Local homozygosity enables RTG recombination in interspecies hybrids

To shed light on how extremely high heterozygosity shapes the RTG recombination landscape, we sequenced the genomes of evolved clones (*n=*53 T6, *n=*5 T12, *n=*4 T24) and non-evolved clones (n=28 T0) derived from three interspecies *S. paradoxus/S. cerevisiae* hybrids (**Table S9**). Genome-wide RTG recombination was less efficient in the interspecies hybrids compared to intraspecies hybrids, generating fewer and smaller LOH (**Fig. 3A-B, Fig S2, Table S6**). Nevertheless, 22 RTG isolates out of 34 had at least one additional recombination event, beside the one selected on chromosome II, and one evolved clone, derived from the SpFE/ScNA hybrid, carried LOH on 3 different chromosomes and spanning 9% of its genome (**Fig. 3B**). This suggested that highly recombined RTG clones do arise in interspecies hybrid populations, but at very low frequencies. To test this possibility, we constructed a SpNA/ScNA hybrid carrying a second unlinked selectable heterozygous marker (*CAN1)*, which enabled us to screen for additional, independent LOH events. Genome sequencing of 4 RTG-clones (T6) selected for loss of *URA3* and *CAN1* revealed clones with multiple LOHs, supporting the scenario that RTG can reshuffle also highly heterozygotic genomes but does so only rarely (**Fig. S7**).

We probed the role of the mismatch repair machinery in suppressing RTG recombination by sequencing SpNA/ScNA evolved and control clones lacking *MSH2* and found ∼1000-fold increase in the median fraction of genome in LOH (2.10 x 10^−4^ vs 1.58 x 10^−7^ in the wild-type). Indeed, LOH formation is hampered by heterozygosity and the inactivation of the mismatch-repair system can mitigate this effect. LOH breakpoints coincided with low local heterozygosity, underscoring that islands of local sequence homology facilitate RTG recombination also in highly heterozygotic hybrids (**Fig. 3C, Fig. S4, Table S7**). Although the scarcity of such islands in highly heterozygous hybrids reduced RTG recombination, the impact of even a single RTG cycle on LOH formation was truly massive compared to the conventional vegetative growth, both in interspecies and intraspecies hybrids (**Fig. S5B**). Interspecies RTG clones were characterised by remarkably stable genomes (**Fig. S5A**), with the only exception of a clone that carried an inversion underlying a complex CNV (**Fig. S7C**). Overall, our results showed that RTG recombination in interspecies hybrids is strongly influenced by heterozygosity, and that inactivation of the mismatch repair machinery mitigates the effect of high sequence divergence.

### Mapping quantitative traits in a sterile hybrid

We probed whether recombination induced by return-to-growth can generate beneficial allelic combinations by measuring the fitness (population doubling time and yield) of 125 ScMA/ScNA recombined RTG clones and 3 T0 samples as controls across 82 environments **(Table S5)**. The fitness of a subset of RTG clones was often inferior to that of the non-evolved hybrid across a wide range of environments. Nevertheless, all RTG recombinants were fitter that their parent in at least some niches and a few were broadly superior, showing that return-to-growth can generate new beneficial allelic combinations capable of driving local adaptation **(Fig. 4A and Fig. S8)**.

**Figure 4.**
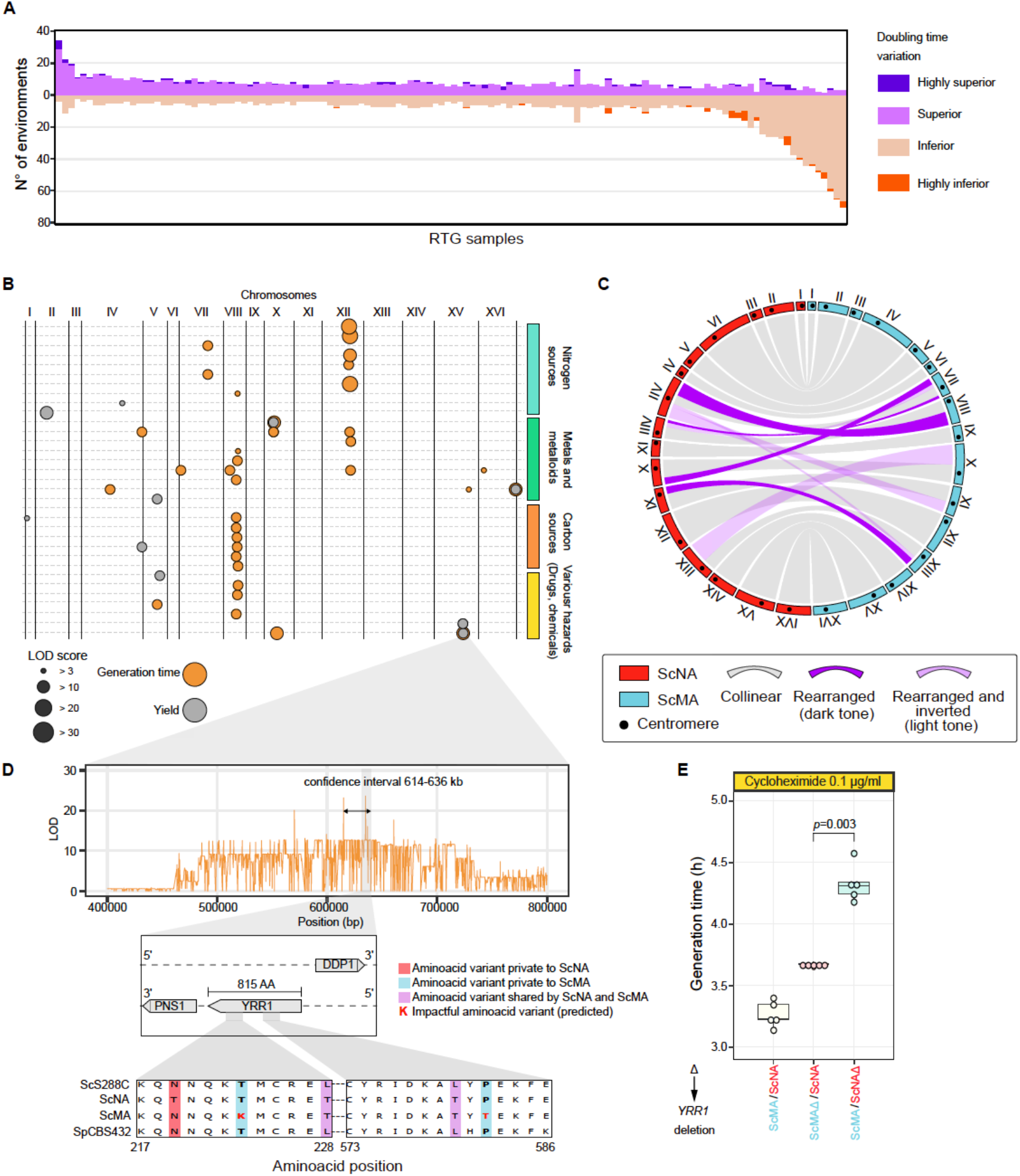
Fitness diversification of RTG clones and QTL mapping. (**A**) Extent of fitness variation upon RTG in the ScMA/ScNA dataset. y-axis shows the number of environments in which the cell doubling time of the RTG clone is superior or inferior to that of the ScMA/ScNA parent hybrid. (**B**) QTLs mapped across the environments. Only conditions where QTLs were detected are reported. (**C**) Circular plot representing the rearrangements between the two subgenomes of the ScMA/ScNA hybrid. Regions are considered inverted if they do not resemble the ancestral centromere-telomere orientation. (**D**) Linkage scan for growth in media containing cycloheximide. Zoom-in on chromosome XV QTL and the two highly conserved regions of *YRR1* with two aminoacids substitutions predicted to be deleterious. (**E**) Boxplot of doubling times in presence of cycloheximide for the parental ScMA/ScNA strains (5 replicates) and the two hemizygous strains (5 replicates). The deletion of either allele reduces growth in cycloheximide compared to the WT, underlying *YRR1* haploinsufficiency in this background. The deletion of the ScNA allele is significantly worse (one-tailed Wilcoxon rank-sum test *p*-value = 0.003) than the ScMA.

Since hybrid sterility precludes standard linkage or association analysis, we next explored whether recombination induced by RTG provides a method for mapping causative genetic variants across lineages that are reproductively isolated post-zygotically. Recombinants that were homozygous for the Malaysian alleles at the *LYS2* locus, suffered a severe fitness decline in 4 environments and became superior in 2 environments. While we could not map the cause of this fitness variation more precisely, we connected fitness variability in 33 environments to 45 quantitative trait loci (QTLs) **(Fig. 4B)**. We found the doubling time in 13 environments to be linked to CNVs resulting from the recombination between non-collinear segments of the North American chromosome VII and the Malaysian chromosome VIII **(Fig. 4B, 4C, Fig. S8)**. Recombinants missing the right arm of Malaysian chromosome VIII (*n*=7) grew slower in all the 13 environments, while those missing the corresponding North American (*n*=2) chromosome arm grew faster or equally fast as non-recombinant samples **(Fig. S8)**.

Next, we asked whether QTL mapping by RTG can be sufficiently resolved to generate a biological understanding of the genetic variants underlying the traits of sterile hybrids. First, we focused on an arsenic resistance QTL located just before the subtelomere of the right arm of chromosome XVI **(Fig. 4B and Fig. S9)**, a region that harbours the *ARR* locus, a gene cluster controlling arsenic exclusion from the cell. This locus is absent in the ScMA subgenome (*11*) and recombinants not inheriting the *ARR* locus were highly sensitive to arsenic (**Fig. S9)**. Then, we explored a major QTL on chromosome XV associated to fitness in presence of the antifungal drug cycloheximide and found near its peak the transcription factor *YRR1* that mediates drug resistance. Since the Malaysian *YRR1* contains two aminoacid substitutions predicted to be deleterious **(Fig. 4D, Table S8)**, we tested whether *YRR1* drives variation in cycloheximide resistance by reciprocal hemizygosity. We found hemizygous strains carrying the Malaysian *YRR1* to be less fit than those carrying the North American allele **(Fig. 4E)**, demonstrating that RTG can generate a genetic understanding of the traits of sterile hybrids.

## Discussion

We evolved 20 diploid yeast genetic backgrounds with varying levels of sterility through RTG, challenging the dogma that sterile hybrids are evolutionary dead ends. We showed that aborting meiosis after its initiation and returning cells to mitotic growth allowed sterile hybrids to overcome reproductive barriers due to mutational defects in late meiosis, structural differences between subgenomes and extreme levels of heterozygosity, and to produce viable recombinant fit clones. These mechanisms constitute some of the post-zygotic reproductive barriers that most commonly underlie speciation in nature, in yeast and other species (*3*). The capacity of one RTG process to drive recombination was substantially lower in interspecies hybrids due to their high heterozygosity and we showed that this barrier to RTG recombination originated in the mismatch repair system. However, abolishing the mismatch repair by removing the mismatch binding protein Msh2 promoted RTG recombination mirroring the increased meiotic recombination observed in gametes of a similar interspecies hybrid (*29*).

The LOH regions generated by recombination between hybrid subgenomes can have profound evolutionary consequences. We have previously shown that the homozygous blocks produced by LOH can mediate meiotic recombination between highly diverged subgenomes, which in turn can rescue hybrid fertility and initiate interspecies introgressions (*30*). Here, we showed that RTG-induced LOHs can make the mating type locus homozygotic, and thereby give rise to mating proficient diploid hybrids. This provides an obvious direct route to polyploidization, and such polyploidization can restore the fertility of sterile hybrids by whole genome duplication (*31, 32*). This RTG driven polyploidization is similar to what is observed in plants, where polyploidization can result from the mating of endoreplicated gametes with an unreduced genome content (*33*), and may help to explain the abundance of yeast polyploids in nature (*34*). LOH regions might be selected by adaptation under specific selective regimes (*35*-*38*) but may also be constrained by incompatibility between allele pairs located in different subgenomes (*39*). Furthermore, the maintenance of a stable diploid state in RTG clones might promote unique evolutionary trajectories, such as the ability to tolerate large CNVs that would otherwise kill haploid gametes. In concert, the impact of RTG recombination on yeast natural evolution has the potential to be quite substantial, and this is underscored by that is arguably quite common, being induced simply by a brief starvation that is followed by cells encountering again nutrients, which must be a occurrence in fluctuating wild habitats (*40*). Whether the role of RTG recombination also extends across broader swaths of the tree of life is unknown, but yeasts separated by hundreds of millions years of evolution from *S. cerevisiae* experienced LOH with a pattern compatible with the RTG signature without a specific tailored protocol (*41, 42*). We see no evident mechanistic reasons for why the RTG process would not extend also across more distantly related sexual organisms, in which meiosis is induced by starvation.

Deviations from the paradigm of sexual reproduction have been repeatedly reported across the eukaryotic tree of life. A meiotic-gene mutant that skips the second meiotic division and produces gametes with unreduced genomic content and complementary recombined genomes was reported in *Arabidopsis thaliana* (*43*). Furthermore, other organisms such as *Candida albicans* (*44*) or the rotifer *Adineta vaga* (*45*) experience genetic recombination without a conventional sexual cycle. The recent characterization of the lifecycle of more than 1000 *S. cerevisiae* strains revealed an independent loss of sexual reproduction in several lineages (*7*). Population genomics data revealed pervasive genome-wide signatures of historical LOHs (*46*) suggesting that RTG might have contributed to the genome evolution of sterile strains. The access to this non-conventional sexual cycle provides a powerful alternative path for genome evolution that breaks the paradigm of sterile hybrids as evolutionary dead ends.

## Supporting information

main_text_figure

phenotypes data

## Authors contributions

A.N. and G.L. conceived the project; S.Mo., L.T., J.W., A.N., G.L., designed the experiments; S.Mo., L.T., A.L., A.I., B.B., N.Š., M.D.C., J-X. Y., M.J. D., A.L., S.L, S.S., R.L., S.Ma., S.S., K.P., performed and analysed the experiments; B.A., performed the sequencing, J.L.L., S.D., J.W., A.N., G.L., contributed with resources and reagents; S.Mo., J.W., A.N., G.L. supervised the project; G.L. coordinated the project; S.Mo. and G.L. wrote the paper with input from L.T., A.N., J.W.

## Data and materials availability

the genome sequences generated in this study are available at Sequence Read Archive (SRA), NCBI under the accession codes: SUB8635978. The phenotype data are available within the supplementary information files. All the strains generated and stored in this work are freely available upon request.

## List of Supplementary Materials

Materials and Methods

References (1 – 20)

Table S1 – S13

Fig. S1 – S9

