## Supplementary material for "Aborting meiosis overcomes hybrid sterility": main_text_figure

### Supplementary Materials

#### Materials and Methods

##### Construction of hybrid yeast strains

All the *Saccharomyces cerevisiae* (Sc) and *Saccharomyces paradoxus* (Sp) strains constructed and used in this study are reported in **Supplementary Table 1**. The diploid hybrid ScS288C/ScSK1 *ndt80Δ* was generated by mating the heterothallic (*ho*) ScS288C and ScSK1 commonly used laboratory strains deleted for *NDT80* (*ndt80::KanMX*) and diploid complementing for histidine auxotrophy were selected. The haploid parental heterothallic (*ho::HygMX*) strains ScNA, ScWA, ScWE, ScSA, ScMA used for generating the intraspecies and interspecies hybrids were previously described (*1*, *2*). In each Sc *MATa* and *MATα* haploid background, the native *URA3* on chromosome V was deleted (*ura3::KanMX*). Subsequently, the *URA3* gene was inserted at the *LYS2* locus in the *MATα* haploid background (*MATα, ura3::KanMX, lys2::URA3*). For generating the intraspecies hybrids, *MATa* cells and *MATα* cells of the two Sc parental species were mated and prototroph diploids (*MATa/MATα, ura3::KanMX/ura3::KanMX, LYS2/lys2::URA3*) were selected on minimal media lacking both lysine and uracil. The haploid parental heterothallic (*ho:HygMX*) strains SpEU, SpFE, SpNA were generated for this work following the genetic engineering scheme explained above for *MATa* and *MATα* strains. For generating the interspecies hybrids, *MATα* cells and *MATa* cells from the Sc and Sp parental species were mated and prototroph diploids (*MATa/MATα, ura3::KanMX/ura3::KanMX, LYS2/lys2::URA3*) were selected on minimal media lacking both lysine and uracil. The ScNA/SpNA *ndt80* and *msh2* diploid mutants were generated by deleting each gene in the respective Sc or Sp haploid background (*ndt80::NatMX* or *msh2::NatMX*) using the lithium acetate/PEG transformation protocol. The haploid strains where mated and diploid prototrophs (*MATa/MATα, ura3::KanMX/ura3::KanMX, LYS2/lys2::URA3, msh2::NatMX/msh2::NatMX*) or (*MATa/MATα, ura3::KanMX/ura3::KanMX, LYS2/lys2::URA3, ndt80::NatMX/ndt80::NatMX*) were selected on minimal media lacking both lysine and uracil. The *CAN1* deletion in the SpNA background was obtained using the lithium acetate/PEG transformation protocol and the diploid hybrid was obtained by mating with ScNA (*MATa/MATα, ura3::KanMX/ura3::KanMX, LYS2/lys2::URA3, can1:NatMX/CAN1*). All the deletions made were verified by polymerase chain reaction (PCR). All the PCR primers used in this study are reported in **Supplementary Table 12**.

##### Selection of clones before and after RTG

Each hybrid was patched from the -80 °C glycerol stock on YPD solid media (1% yeast extract, 2% peptone, 2% dextrose, and 2% agar) and incubated overnight at 30 °C. The following day the strain was streaked to minimal solid media not supplemented with uracil and the plate was incubated at 30 °C for 48 hours. Five single colonies of the hybrid strain were taken and inoculated separately in 10 mL of YPEG pre-sporulation medium (1% yeast extract, 2% peptone, 3% ethanol, and 3% glycerol) for 15 hours at 30 °C with shaking at 220 rpm. Each pre-sporulation culture was washed twice with sterile water and resuspended in 2% potassium acetate to reach an $OD_{600} = 0.5$ in a 250-mL flask. One mL was immediately collected from the starving culture to generate the T0 sample (before meiosis). The flasks were incubated at 23 °C, with shaking at 220 rpm, and after 6 hours of incubation the T6 sample were taken. The T0 and T6 samples were washed twice with 1 mL of YPD and incubated in 1 mL of nutrient rich YPD for 18 hours at 30 °C without shaking, thereby aborting meiosis in the T6 samples and returning cells to growth (RTG). The following day the YPD liquid cultures were vortexed and, depending on the strain and the density of the culture, 20 to 200 µL of 10-fold diluted culture were plated on minimal medium containing 1 mg/mL of 5-fluoroorotic acid (5-FOA) and spread with glass beads (*3*). In parallel, cells from the same YPD liquid culture were serially diluted up to $10^{-5}$ and spotted on YPD plates in at least two replicates for each biological replicate. The YPD and 5-FOA plates were then incubated at 30 °C for 48 hours. After that, colonies growing on 5-FOA (and therefore having lost their *URA3* marker through loss-of-heterozygosity) and YPD plates were counted and used to infer CFU/mL in each condition. RTG clones from the S288C/SK1 hybrids were obtained using the mother-daughter protocol as previously described (*4*).

**Meiosis dynamics and spore viability**

The meiosis of diploid hybrids engineered with the *LYS2*/*lys2::URA3* system was induced with the same protocol used for RTG but the flasks were kept at 23 °C with shaking at 220 rpm and monitored for the formation of viable spores (germinated gametes). In order to analyse the sporulation efficiency (fraction of cells having passed through meiosis), at least 200 cells were counted for each sample and estimated the percentage of those that had sporulated (formed dyads, triads or tetrads of spores). To estimate spore viability (gametes that are viable and capable of germinating), complete tetrads were dissected and deposited spores on YPD plates where they were allowed to germinate and grow for four days at 30 °C. The spore viability was calculated as the fraction of spores that had germinated and formed visible colonies. To monitor meiotic progression by DAPI staining 2 mL of T0, T6 and T12 culture were collected. Each of the 2 mL samples were put in a separate tube with 5 mL of EtOH 70% and stored at -20 °C. Frozen samples were washed twice with 1 mL of sterile water, resuspended in 200 µL of water and stained with 2 µL of DAPI (4′,6-diamidino-2-phenylindole) for 30 minutes in the dark. Cells were counted using a fluorescence microscope.

**RTG selection by *URA3*-loss assay and *URA3*/*CAN1*-loss assay**

We designed the 5-FOA assay to detect the increase of recombination upon RTG at the heteroallelic locus *LYS2/URA3* on chromosome II. We deleted the copy of *URA3* from its native location on chromosome V in all the haploid parental strains. We replaced one *LYS2* allele on chromosome II with one copy of *URA3* in one of the two parents used to generate each diploid hybrid, and performed the assay as already reported in the literature (*5*). The growth in YPD for 18 hours ensured that all the cells that were in the early phase of meiosis performed RTG. We used the same incubation time in liquid YPD for the sporulating cultures (T6) and the controls (T0) to take into account LOH occurring during the growth in YPD. We confirmed by whole-genome-sequencing of single clones that *de novo* mutations and aneuploidies were not impacting the heteroallelic assay, and all events that lead to *URA3* loss were LOHs. Therefore, we concluded that T0 cells growing on 5-FOA were due to mitotic LOH whereas T6 cells had a composite effect of mitotic and RTG-induced recombination. Sequencing of single clones supported this scenario, and T6 RTGs had a single LOH event on chromosome II (mitotic recombination or low efficiency RTG), or additional LOHs (RTG) in the genome. This difference must be carefully considered also for interspecies hybrids, although in those backgrounds the lack of additional LOH events can result from anti-recombination mechanism due to the high sequence divergence. We expanded this protocol to introduce an additional selection step as already reported in the work by Coelho *et al*. in which multiple selection steps were used (*6*) and also based on an early work in which RTG cells had recombination at unlinked genetic markers (*7*). We introduced an additional selection marker by deleting one copy of *CAN1* on chromosome V in one of the two parents of the hybrid tested, so that two independent LOHs could be selected for, one on chromosome II, at the *LYS2/URA3* locus, and one on chromosome V, at the *CAN1/CAN1::NatMX* locus. Briefly, after 18 hours of YPD incubation RTG cells were diluted 1:10 to an OD of ~0.5 in SD media supplemented with canavanine and grown for 10 hours. Finally, cells were plated on 5-FOA plates following the protocol described in the paragraph "Selection of clones before and after RTG" in the Materials and Methods section. The canavanine liquid incubation enriches for cells bearing LOH at the *CAN1* locus and the passage on 5-FOA plates selects for cells carrying LOH at the *LYS2/URA3* locus.

**Sequencing data analysis**

All the samples were sequenced with Illumina paired-end technology at the NGS platform of Institut Curie according to the manufacturer's standard protocols. Short-read sequencing data were processed by means of the MuLoYDH pipeline using default parameters and the assemblies/annotations embedded in MuLoYDH (*5*). All the data-sets consisted of the hybrids evolved under the RTG protocol and analysed against the corresponding parent hybrid before RTG as control samples **(Supplementary Table 9)**. The pipeline required as input: (1) a data set of short-read sequencing experiments from yeast diploid hybrids, and (2) the two parental genomes which were used to produce the hybrids in FASTA format as well as the corresponding genome feature annotations in the "general feature format" (GFF). The availability of reference-quality genome assemblies for all the parents used to generate the panel of hybrids, allowed a highly accurate tracking of the mutational landscape. Reads from sequencing data were mapped against the assemblies of the two parental genomes separately (standard mappings) and against the union of the two aforementioned assemblies (namely a multi-FASTA obtained by concatenating the two original assemblies) to produce the competitive mappings. In the latter case, reads from parent 1 were expected to map to the assembly of parent 1 on the basis of the presence of single-nucleotide markers. Conversely, reads from parent 2 were expected to map to the assembly of parent 2.

Standard mappings were used to determine the presence of CNVs. The latter were also exploited to discriminate LOHs due to recombination from those resulting by deletion of one parental allele. The markers between the parental assemblies were determined by the NUCmer algorithm (v 4.0.0 beta) and were exploited to map LOH segments. Markers were genotyped from standard mappings. Marker positions characterized by nonmatching genotype or alternate allele were filtered out, as well as multiallelic sites, whereas those lying in subtelomeric and telomeric regions were masked. Remarkably, MuLoYDH provided LOH calls without filtering out small events using an arbitrary threshold based on the number of supporting markers. Indeed, we are able to detect events supported by a single marker and we previously demonstrated that such events are genuine LOH (*5*). Stretches of consecutive markers showing homozygous genotypes were grouped in LOH regions. The genomic coordinates of each LOH event were determined using both the "first/last" coordinates and the "start/end" coordinates. First/last coordinates were determined using the coordinates of the first and the last markers of the event. Start/end coordinates were calculated using the average coordinate of the first (last) marker and the last (first) marker of the adjacent event. LOH regions were annotated as terminal/interstitial. Interstitial LOHs were defined as homozygous segments flanked on both sides by heterozygous markers. Terminal LOHs were defined as homozygous regions extended to the end of the chromosomal arm, i.e. the last non-masked marker.

Due to low coverage that did not match the standard requested by MuLo, we excluded two WT RTG derived from ScS288C/ScSK1. We also removed one sample that went through two cycle of RTG in order to have only samples that performed one RTG cycle in WT and *ndt80Δ*.

*De novo* single-nucleotide variants and indels were determined from both competitive and standard mappings. Competitive mapping allowed for direct variant phasing in heterozygous regions. Variant calling from competitive mapping was performed setting ploidy = 1 in heterozygous regions and ploidy = 2 in LOH blocks. Regions characterized by reads with low mapping quality (MAPQ < 5 in the competitive mapping) were assessed from standard mapping using arbitrarily the assembly from parent 1. All the *de novo* variants detected were checked by visual inspection using IGV (*8*).

**Testing for association between LOH breakpoints and Spo11p-induced DNA double-strand breaks (DSB) or recombination hotspots**

The association between LOH breakpoints and DSB/recombination hotspots was tested using the *regioneR* package by means of the function *overlapPermTest*, setting the parameters: ntimes = 10000, and alternative = "greater". The *fasta* suite (*9*) was used to convert the coordinates of the hotspots regions detected in the original reference genomes (SGD_R62-1-1_20090218 and SGD_R58-1-1_20080305 for the hotspots reported by Pan *et al*. (*10*) and Mancera *et al*. (*11*) respectively) to the corresponding coordinates of the hybrid subgenome engineered with the *lys2::URA3* selection system.

**Genome content analysis of polyploid strains**

The DNA content of the mating proficient diploid RTGs was analysed upon mating with tester strains using a propidium iodide (PI) staining assay. Cells were first pulled out from glycerol stocks on YPD solid media and incubated overnight at 30 °C. The following day a small portion of each patch was taken with a pipette tip, transferred in 1 mL of liquid YPD and incubated overnight at 30 °C. Then, cells were washed with water, resuspended in 1 mL of cold 70% ethanol and fixed overnight at 4 °C. Finally, the samples were washed twice with phosphate-buffered saline (PBS), and 100 µL of each sample was resuspended in 900 µL of staining solution (15 µM PI, 100 µg/mL RNase A, 0.1% v/v Triton-X, in PBS) and incubated for 3 hours at 37 °C in the dark. Ten thousand cells for each sample were analysed on a FACS-Calibur flow cytometer. Cells were excited at 488 nm and fluorescence was collected with a FL2-A filter. The data were analysed using the R packages flowCore (*12*) and flowViz (*13*), and were plotted with ggplot2 (R version 3.6.1).

**Long reads sequencing and structural variant analysis**

Yeast cells were grown overnight in liquid YPD media. Genomic DNA was extracted using Qiagen Genomic-Tips 100/G according to the manufacturer's instructions. The MINION sequencing library was prepared using the SQK-LSK109 sequencing kit according to the manufacturer's protocol. The library was loaded onto a FLO-MIN106 flow cell and sequencing was run for 72 hours. Long read basecalling and scaffolding were performed using the pipeline LRSDAY (v 1.6) (*14*) and the dotplot of chromosome III was generated using mummerplot (*15*). The inversion on chromosome III was detected using sniffles (*16*) implemented within the Varathon framework (https://github.com/yjx1217/Varathon).

**Estimating growth during mitotic reproduction**

All yeast strains were stored at -80 °C in 20% glycerol and cultivated at 30 °C in temperature and humidity-controlled cabinets. Yeast strains were revived from frozen 96-well stocks by robotic transfer (Singer

RoToR; long pins) of a random sub-sample of each thawed population (~50,000 cells) to a Singer PlusPlate in 1536 array format on solid Synthetic Complete (SDC) medium composed of 0.14% Yeast Nitrogen Base (CYN2210, ForMedium), 0.50% $(NH_4)_2SO_4$, 0.077% Complete Supplement Mixture (CSM; DCS0019, ForMedium), 2.0% (w/w) glucose, pH buffered to 5.80 with 1.0% (w/v) succinic acid and 0.6% (w/v) NaOH. In every fourth position, fixed spatial controls (genotype: YPS128, *MATa*/*MATα*) were introduced to account for spatial variation across plates in the subsequent experimental stage. Controls were similarly sub-sampled from a separate 96-well plate and introduced to the pre-culture array (Singer RoTor; long pins). Populations were pre-cultivated for 72 hours at 30 °C. For pre-cultures of nitrogen-limited environments, the background medium was modified to avoid nitrogen storing and later growth on stored nitrogen: CSM was replaced by 20 mg/L uracil (not converted into usable nitrogen metabolites) and $(NH_4)_2SO_4$ was reduced to growth-limiting concentrations (30 mg/L of nitrogen). We cast all solid plates 24 hours prior to use, on a levelled surface, by adding 50 mL of medium in the same upper right corner of the plate. We removed excess liquid by drying plates in a laminar air-flow in a sterile environment. Pre-cultured populations were mitotically expanded until stationary phase (2 million cells; 72 hours), were again subsampled (~50,000 cells; short pins) and transferred to experimental plates, containing the medium of interest (**Supplementary Table 5**). Synthetic grape must (SGM) was prepared as previously described. We tracked population size expansion using the Scan-o-Matic system, version 1.5.7 (https://github.com/Scan-o-Matic/scanomatic) (*17*). Plates were maintained undisturbed and without lids for the duration of the experiment (72 hours) in high-quality desktop scanners (Epson Perfection V800 PHOTO scanners, Epson Corporation, UK) standing inside dark, humid and thermostatic cabinets with intense air circulation. Images were analysed and phenotypes were extracted and normalized against the fourth position controls using Scan-o-Matic. We extracted the normalized, relative population size doubling time, $D_r$, by subtracting the control value for that position, $D_r = \log_2(D) - \log_2(D_{control,local})$. $D_r$ is reported as output data. Growth phenotypes were qualitatively classified as inferior, when $D_r$ normalized to the initial hybrid was between 0.25 and 1, highly inferior when $D_r$ was greater than or equal to 1, superior when $D_r$ was between -0.25 and -1 and highly superior when the $D_r$ was less than or equal to -1.

##### QTLs mapping and phenotype analysis

QTL analysis was performed with the package R/qtl (v1.46-2) (*18*) using mitotic growth data normalized with the growth data of the parental hybrid for 125 RTG clones and 3 T0 samples of the ScNA/ScMA hybrid as the phenotype variable and the genotype of heterozygous markers extracted from the sequence data of the same RTG clones as the genotype variable. We removed markers from the dataset to which the pipeline did not assigned any genotype with the function “drop.nullmarkers”, and we also removed markers for which only one sample was found to be homozygous. We estimated a genome-wide LOD statistical threshold performing 1000 random permutations of both the phenotype and genotype rows and extracting the 95$^{th}$ percentile of LOD values as threshold. The confidence interval of the QTL was calculated using the "lodint" function of R/qtl (*18*).

**Reciprocal hemizygosity assay and functional mutation prediction**

Start to stop *YRR1* gene deletions was engineered in the parental haploid strains CC407 (ScNA) and YGL1027 (ScMA) by replacing the open reading frame with the NatMX cassette. The haploid strain CC407 *yrr1*::NatMX was then crossed with a wild-type haploid YGL1027 to obtain a hemizygous diploid carrying only the MA *YRR1* allele. and the haploid YGL1027 *yrr1*::NatMX strain was crossed with a wild-type haploid CC407 to obtain a hemizygous diploid carrying only the NA *YRR1* allele.

Cells from the wild-type ScMA/ScNA hybrid and the two hybrids reciprocally hemizygous for *YRR1* were pre-grown in a 96 well-plate containing YPD for 16 hours in 5 biological replicates for each strain background. The following day 20 µL of cultures were taken from each well and transferred to another 96 well-plate containing 180 µL of YPD with cycloheximide at a concentration of 0.1 µg/mL. The growth was monitored by measuring OD changes over 72 hours on a Tecan plate reader (infinite F-200 pro). The generation time was extracted from the OD profile using the software PRECOG (*19*) and then plotted and analysed in Rstudio. The prediction of deleterious variants was obtained from the mutfunc suite (*20*).

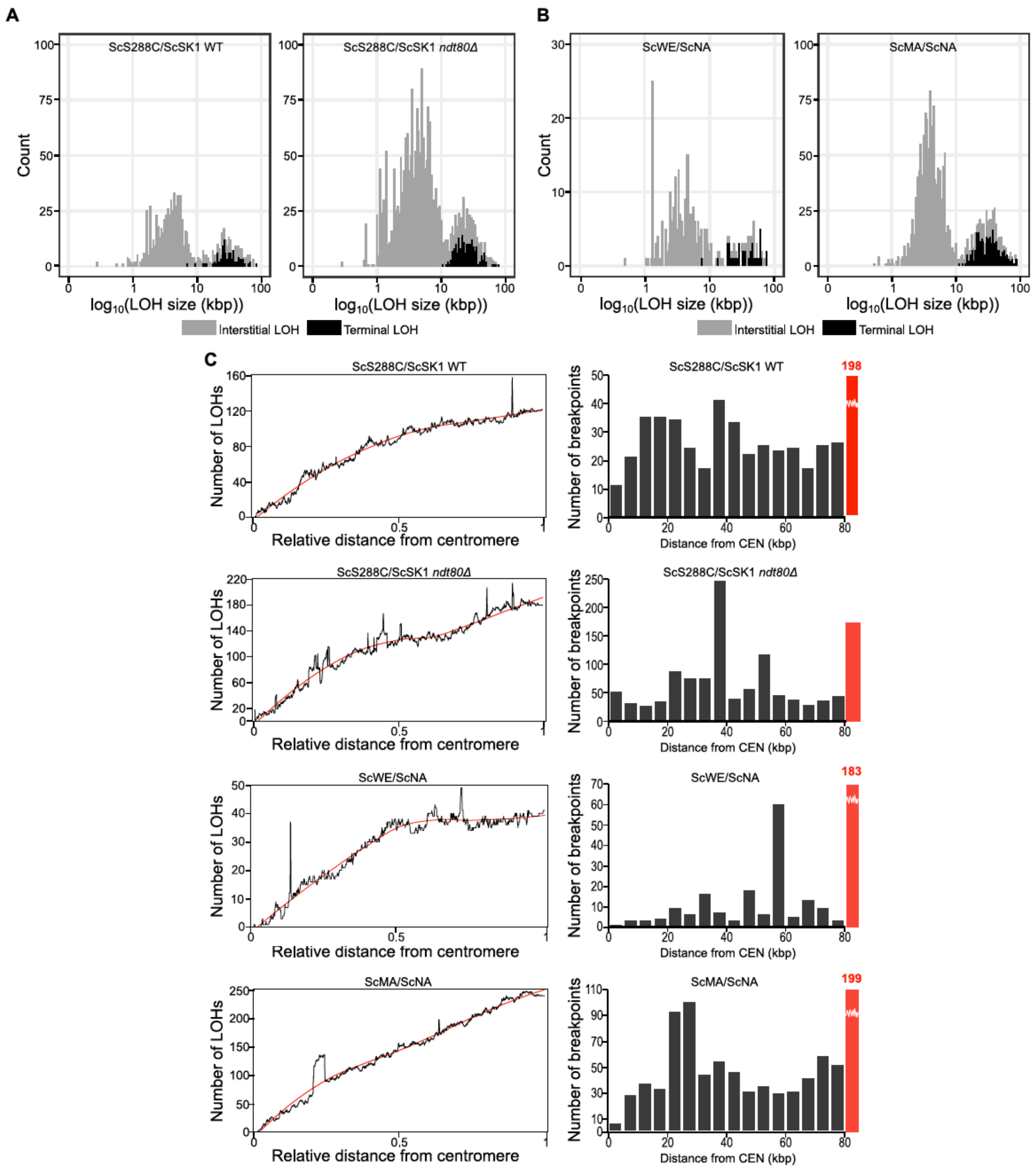


**Supplementary Figure 1. LOH size and chromosomal position.** (**A**) Size distribution of interstitial (grey) and terminal (black) LOH events for the ScS288C/ScSK1 WT and *ndt80*Δ RTGs. The x-axis represents the $\log_{10}$ of the size of the LOH. (**B**) Size distribution of interstitial (grey) and terminal (black) LOH events for the intraspecies (light brown) and interspecies (dark brown) RTGs. The x-axis represents the $\log_{10}$ of the size of the LOH. (**C**) Left panels: number of LOHs versus the relative distance from the centromere. For each background, the number of LOH events spanning 1000 bins of width 0.001 is reported as a black line. The plots represent aggregate analysis for all chromosomes except for events lying on chromosome II that were filtered out since positions of LOH in this chromosome were constrained by the presence of the *LYS2/URA3* system (see text). The smoothing of the data (red line) serves as an eye guide. Right panels: bar plot of the number of breakpoints in a 80 kbp window around all the centromeres (excluding chromosome II) for the intraspecies RTGs with 5 kbp bin size. Red bars represent the whole-genome median. The values exceeding the y-axis upper limit are annotated on top of the corresponding bar.

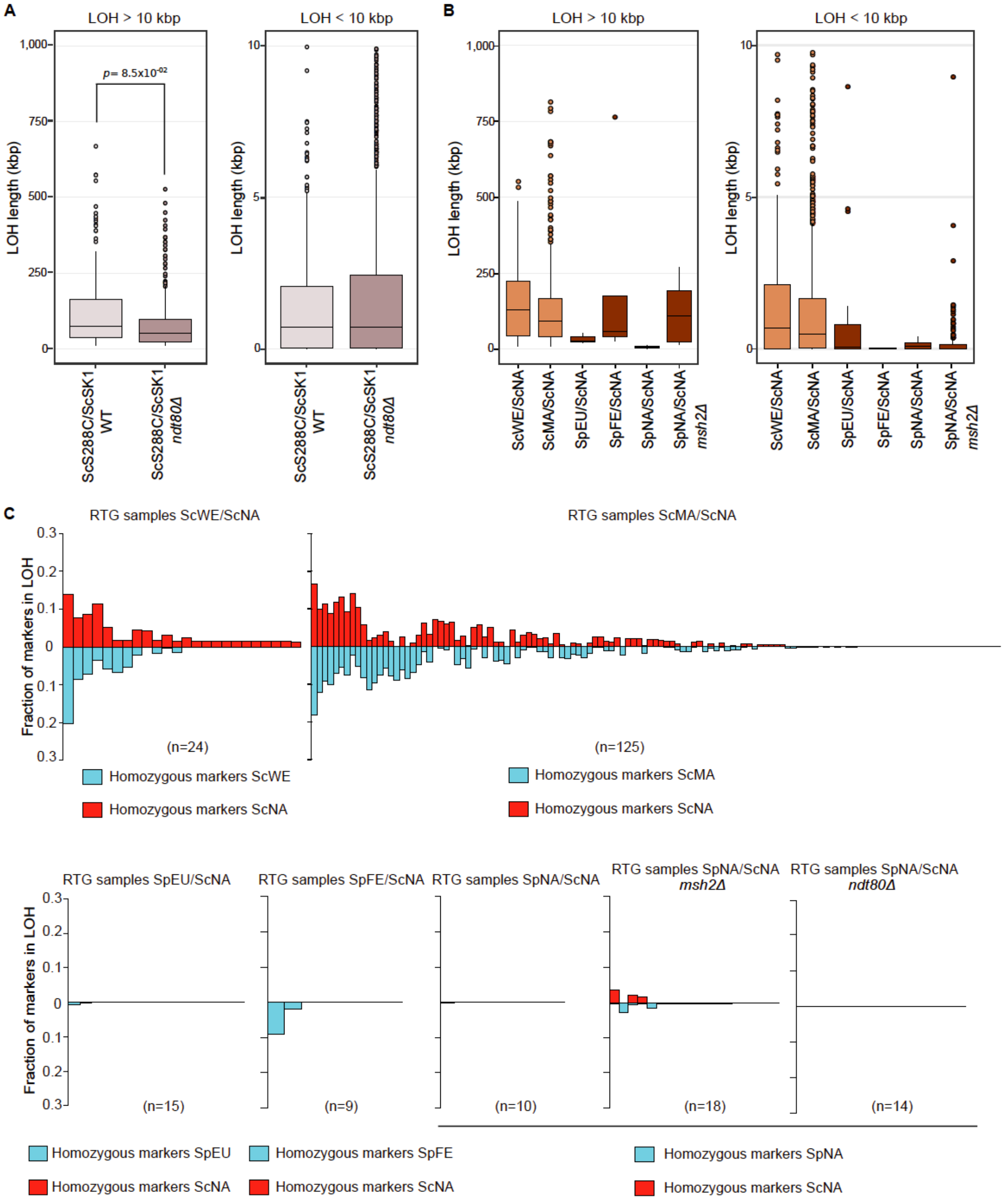


**Supplementary Figure 2. Overview of LOH length and heterozygous/homozygous markers in RTG clones.** (**A**) Boxplots of S288C/SK1 and S288C/SK1 *ndt80Δ* LOHs (left > 10 kbp and right < 10 kbp). The *p*-value refers to two-tailed Wilcoxon rank-sum test. The same test performed only on the mother cells of the *ndt80Δ* and WT populations confirmed this result (one sided Wilcoxon rank-sum test, *p*-value=$2.34 \times 10^{-8}$) (**B**) Boxplots of the LOH (left > 10 kbp and right < 10 kbp) for the intraspecies and interspecies hybrids colored according to figure 2. (**C**) RTG clones from intraspecies hybrids display high heterogeneity in the genome fraction in LOH (upper panels). We did not observe any LOH bias towards one of the two hybrid's subgenomes for intraspecies hybrids (Welch's t-test ScWE/ScNA, p-value = 0.8, Welch's t-test ScMA/ScNA, p-value = 0.74). Interspecies hybrids show a lower magnitude of genome in LOH (lower panels).

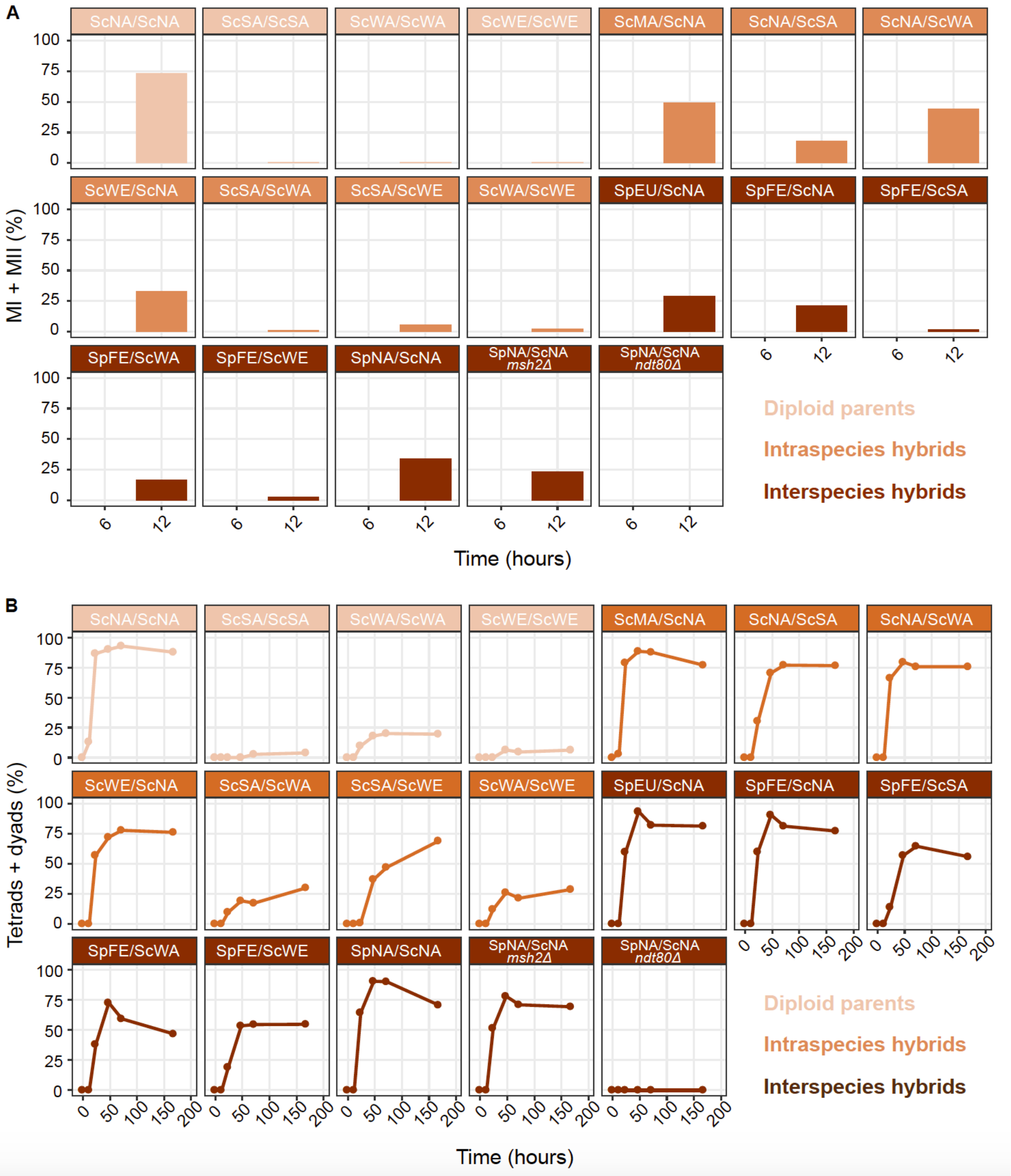


**Supplementary Figure 3. Sporulation dynamics across the strain panel.** (**A**) Meiotic progression measured by DAPI staining as the combined percentage of cells with 2 nuclei (MI) and 4 nuclei (MII). None of the samples had 2 or 4 nuclei at T0 (not reported) and T6. (**B**) Sporulation efficiency measured as the combined percentage of tetrads and dyads observed on the total number of counted cells.

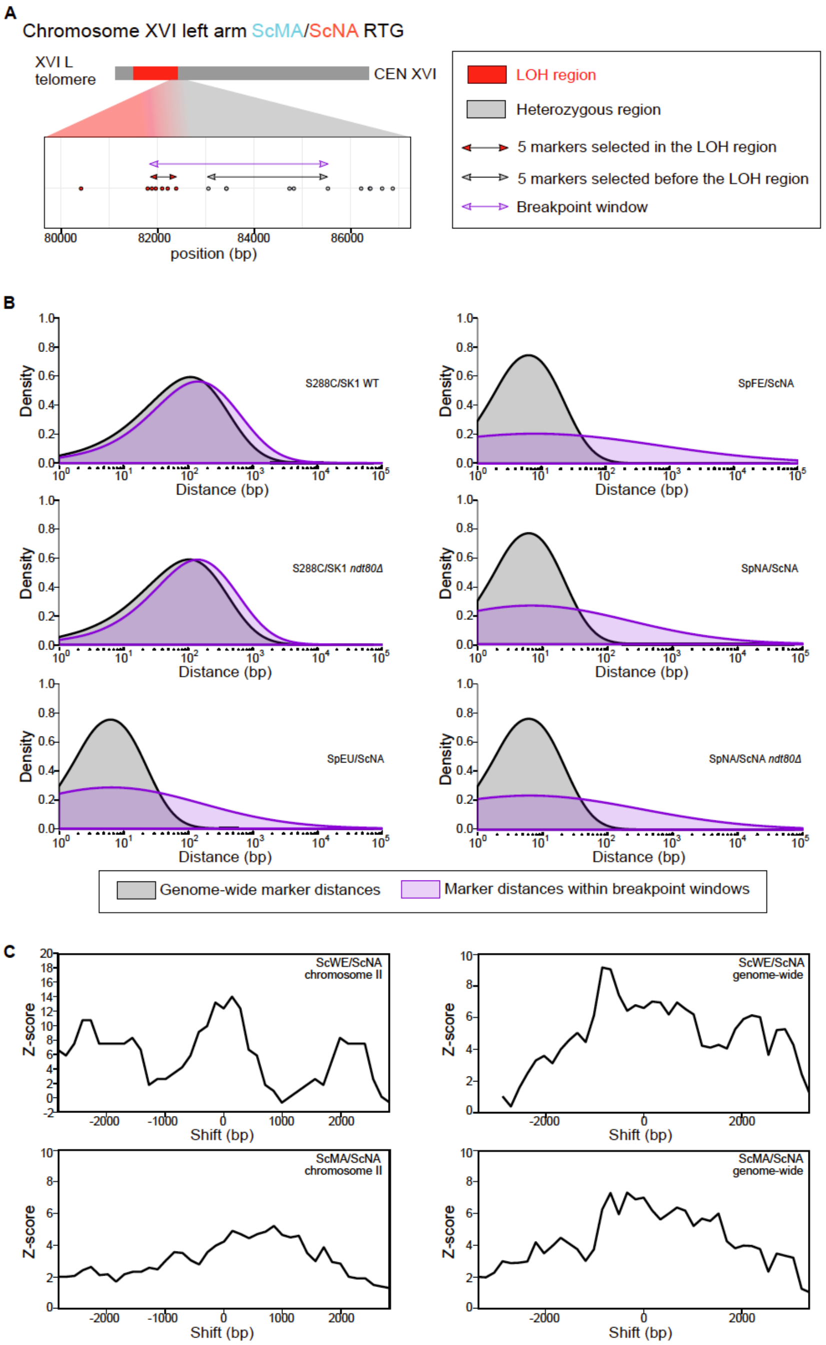


**Supplementary Figure 4. LOH breakpoints are enriched in low heterozygosity regions and nearby meiotic recombination hotspots. (A)** Zoom-in of a LOH breakpoint window that highlights which markers were selected to evaluate the local heterozygosity (see Figure 3 and Figure S4B). LOH breakpoint windows comprise the 5 heterozygous markers and the 5 homozygous markers closer to the breakpoint. **(B)** Distribution of marker distances genome-wide (grey) and in LOH breakpoint windows (purple) across other sequenced hybrids. **(C)** Z-score of the statistical test for LOH breakpoints and recombination hotspots association on chromosome II (left panel) and genome-wide (right panel) for the ScWE/ScNA and ScMA/ScNA evolved samples. The Z-score is plotted as a function of the shift (in bp) of the recombination hotspots regions.

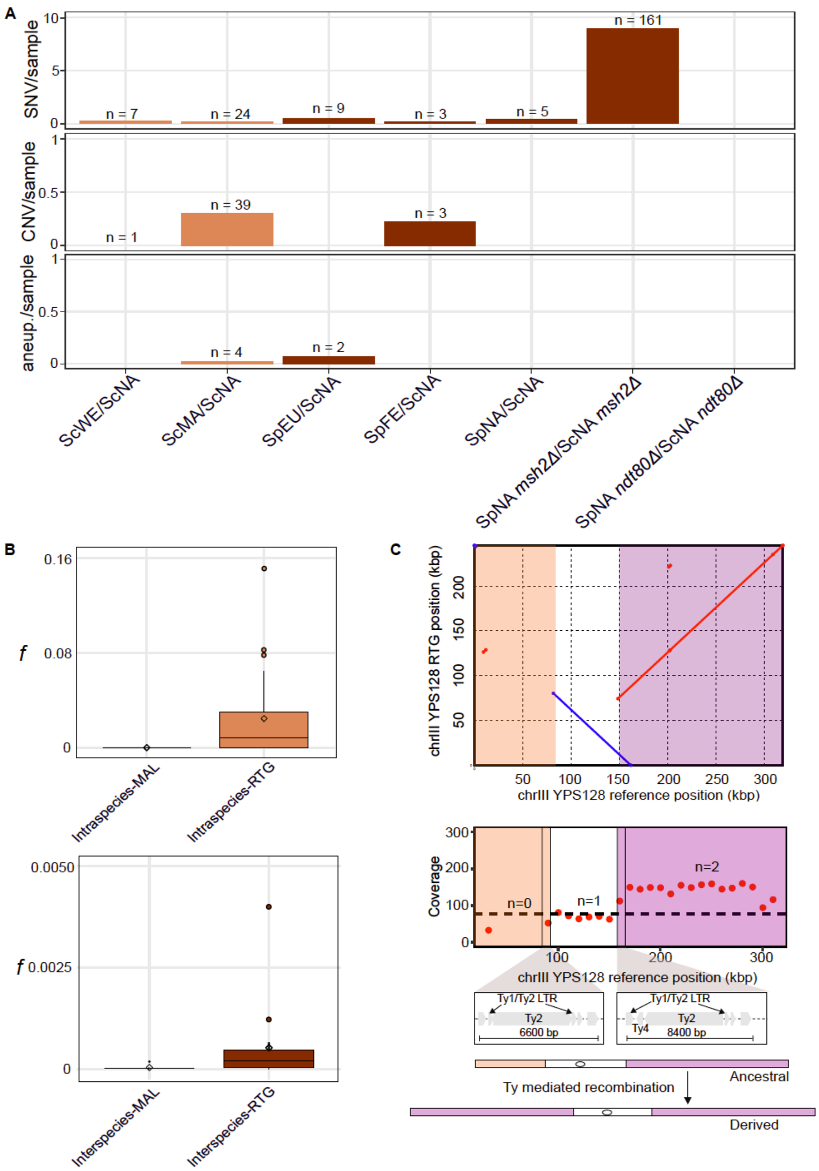


**Supplementary Figure 5. Mutational landscape upon RTG.** (**A**) Bar plots report the number of single nucleotide variants (SNVs), copy number variations (CNVs) and aneuploidies per sample. Genomes of evolved RTG clones are remarkably stable with no significant increase of aneuploidies, single nucleotide variants (SNVs) and copy number variations (CNVs). The increased SNVs in the SpNA/ScNA *msh2Δ* hybrid is expected given its mutator phenotype. Massive CNVs in the ScMA/ScNA samples arise as a result of recombination between non-collinear chromosome arms. (**B**) Top panel: LOH genomic impact on intraspecies ScWE/ScNA hybrids evolved through mutation accumulation lines (MAL) and RTGs; *f* is the fraction of genome in LOH per genome per bottleneck. Events lying on chromosome II of RTG samples were filtered out. Bottom panel: LOH genomic impact for interspecies SpEU/ScWE hybrids evolved through MAL and the SpEU/ScNA RTGs. (**C**) Top panel: dot plot of chromosome III from Nanopore long-read sequencing and *de novo* assembly of an RTG clone which underwent a complex rearrangement resulting in loss of the distal part of the left arm, inversion of the region encompassing the centromere and duplication of part of the right arm. The Illumina short-read coverage analysis support this rearranged structure, which was likely due to a Ty-mediated mechanism (bottom panel).

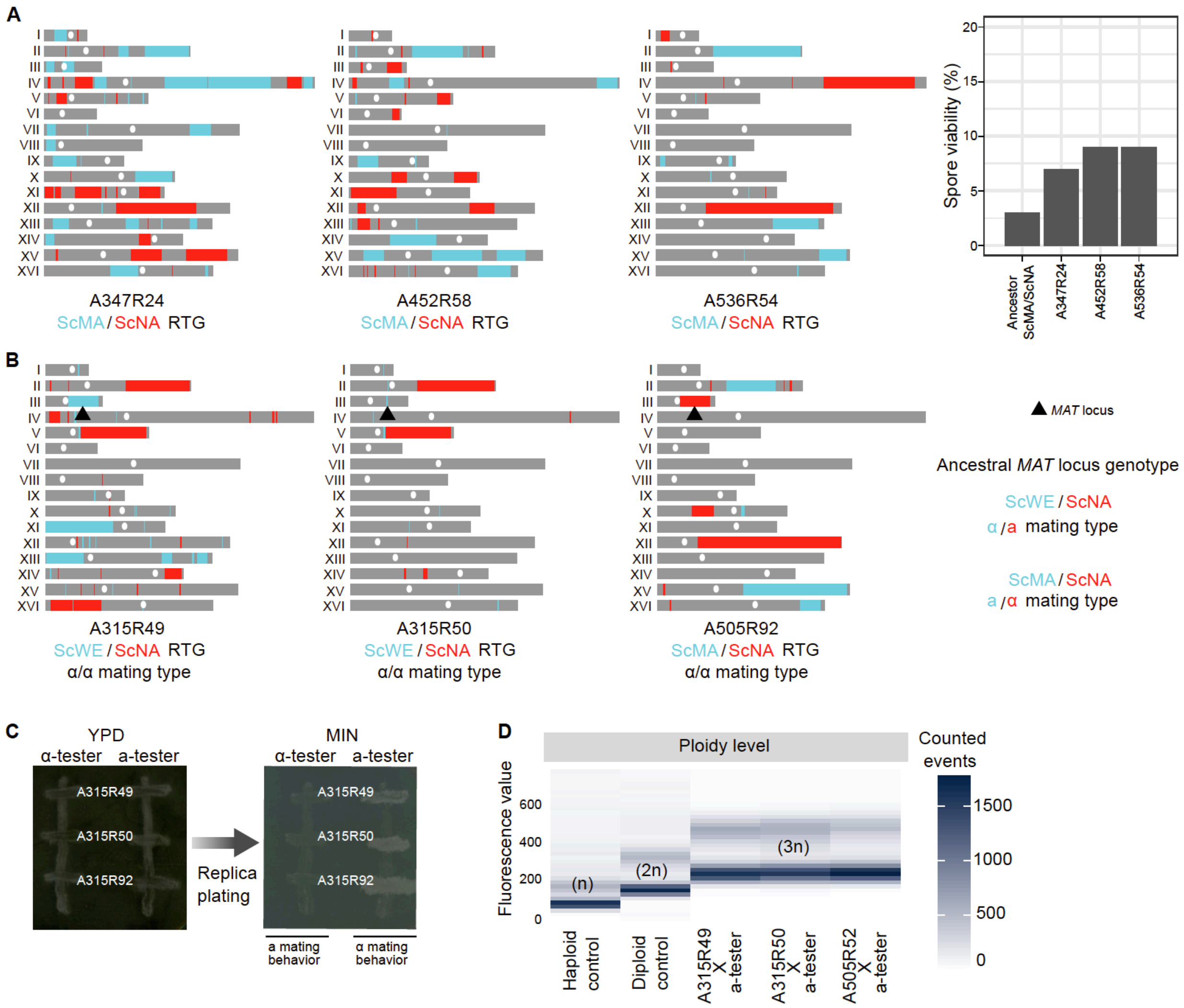


**Supplementary Figure 6. RTG partially rescues hybrid fertility and can lead to polyploidization.** (**A**) LOH landscapes of 3 selected ScMA/ScNA RTGs clones (left) and their respective spore viability compared to the ancestral hybrid (bar plot, right). (**B**) LOH landscape of two ScWE/ScNA RTGs and one ScMA/ScNA RTG with α/α mating type. (**C**) The mating-type test shows that RTG clones with recombination encompassing the *MAT* locus have haploid-like mating behavior and can generate polyploid strains if mated with a partner of the opposite mating-type. (**D**) The genome content of the mated strains has been validated by flow-cytometry.

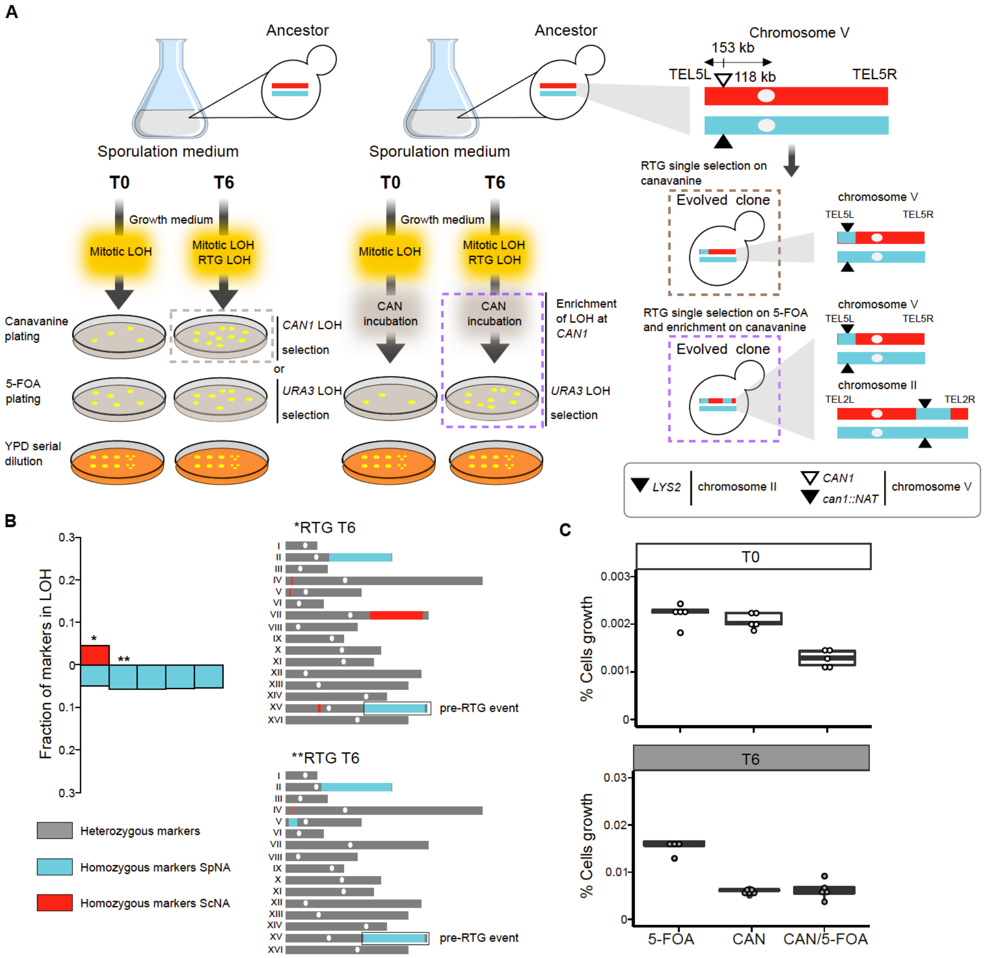


**Supplementary Figure 7. Overview of the double LOH selection approach.** (**A**) Left panel: RTG protocol used for the single selection approach performed either with 5-FOA or canavanine. Central panel: RTG protocol used for the enrichment of a second recombination event when cells are grown on canavanine before plating on 5-FOA. Right panel: sketch of the LOH recombination selected when plating on canavanine (brown box) and the event enriched upon canavanine incubation and plating on 5-FOA (purple box). The *CAN1* gene is deleted in the *S. paradoxus* subgenome so that the two selectable markers promote LOH in the same direction to avoid any genetic incompatibility effect by selecting two LOHs toward different subgenomes. (**B**) Fraction of markers in LOH in the sequenced RTG isolated with the double selection approach and LOH map of two RTGs sequenced. (**C**) Percentage of cells growing on 5-FOA or canavanine plates upon RTG. Three experiments are reported: single selection approach (5-FOA or canavanine) and double selection approach. T0 = no sporulation induction, T6 = 6 hours of sporulation induction. The incubation in canavanine does not kill the cells but allow to grow only those having resistance to the drug thus enriching cells harbouring LOH at *CAN1*. Once the cells are plated if the two events were completely independent the ratio of T6 compared to the T0 would have been expected to be the same and cells with only the LOH on chromosome II deleting *URA3* were expected to be retrieved.

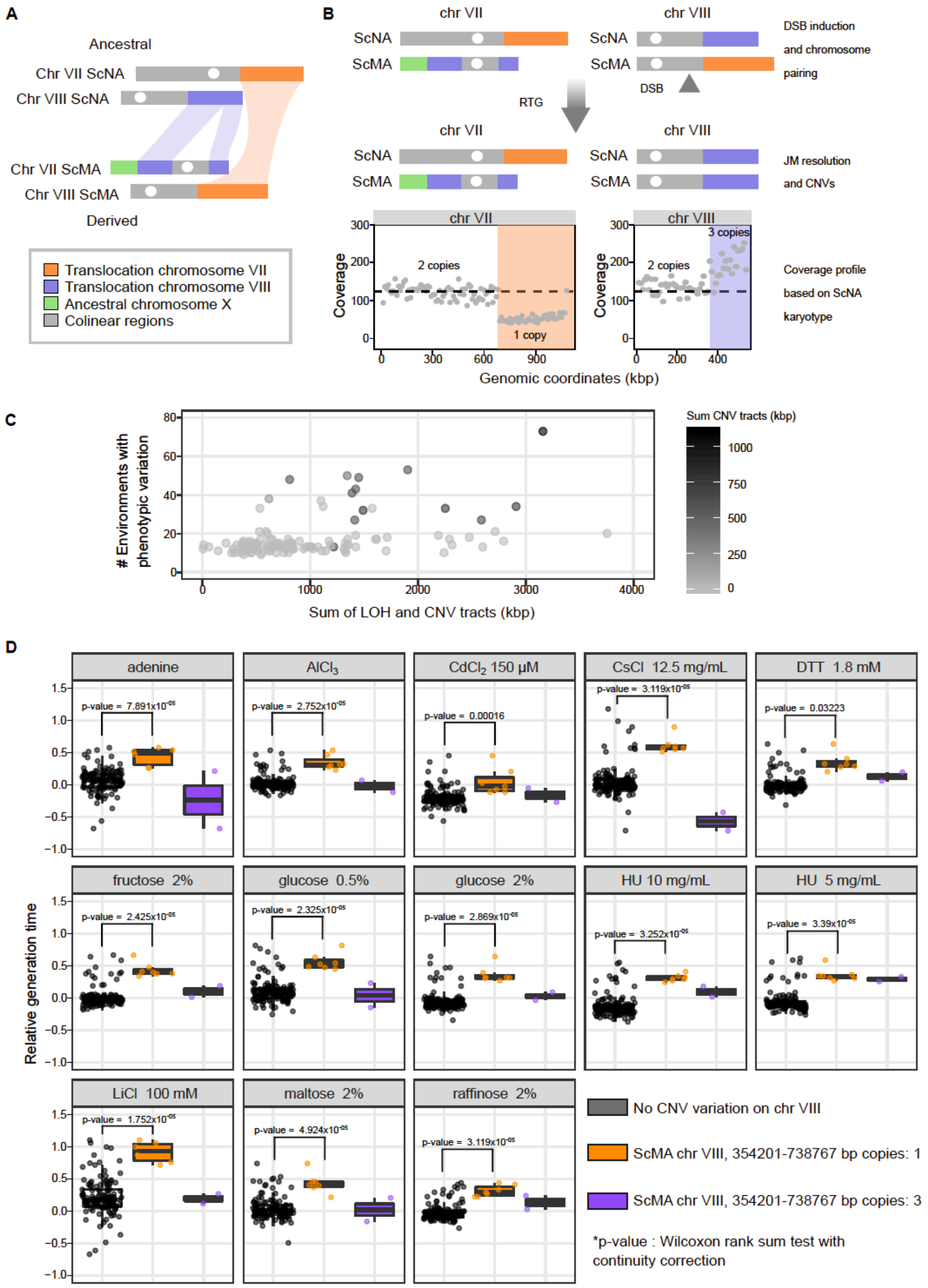


**Supplementary Figure 8. Massive CNVs shape RTGs fitness.** (**A**) Sketch of the non-colinear regions on chromosomes VII and VIII. The term "ancestral" refers to the ScNA karyotype whereas the term "derived" refers to the rearrangements in the ScMA karyotype. (**B**) Massive CNVs result from recombination of non-collinear chromosome arms of the subgenomes of a ScMA/ScNA hybrid. (**C**) Relationship between phenotypic variation and length of LOH and CNV events in the RTG samples; 2 samples were removed due to non-reliable CNV calls. (**D**) Boxplots of the relative generation time of RTGs with no CNV on chromosome VIII (black), RTG that have lost one copy of the arm of chromosome VIII of the ScMA subgenome as a result of recombination (orange), and RTG that have gained one copy of the arm of chromosome VIII of the ScMA subgenome as a result of recombination (purple).

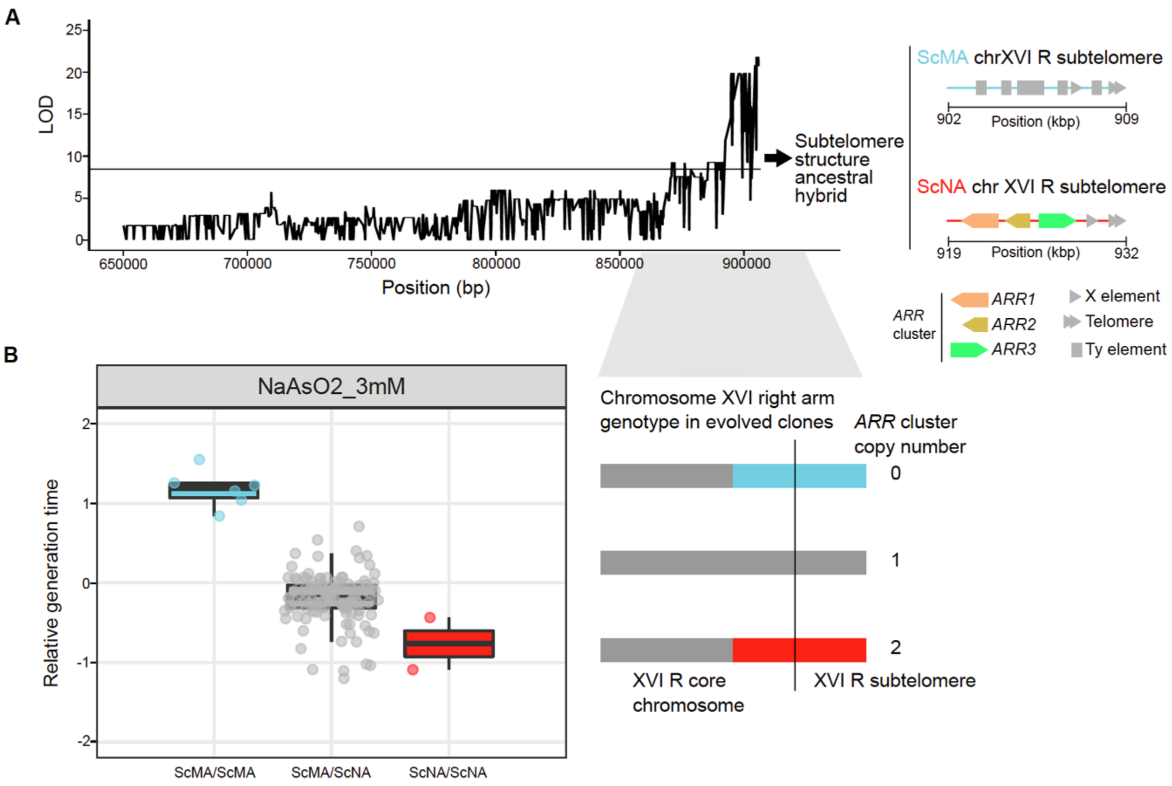


**Supplementary Figure 9. Linkage analysis of arsenite resistance.** (**A**) Plot of the LOD score along the right arm of chromosome XVI. The horizontal line indicates the statistical significance threshold. Right side show a zoom-in on the subtelomere structure of the two parental isolates used in the linkage mapping. The ScMA background lack the arsenite resistance cluster *ARR1-3*. (**B**) Phenotypic variation in arsenite across the ScMA/ScNA RTG samples. The data points in the boxplots are grouped based on the genotype of the last marker before the right subtelomere of chromosome XVI. The right panel shows the different number of *ARR* clusters according to the genotype of RTG clones.
